# Quantitative Optimization of Sensitivity and Specificity in Targeted and Whole-Exome Sequencing Using Reference-Standard DNA Mixtures

**DOI:** 10.64898/2026.01.25.701479

**Authors:** Youngbeen Moon, Chung Hwan Hong, Jong-Kwang Kim, Eun-Kyung Kang, Hye Won Choi, Dong-Won Hwang, Jae-Hwan Ko, Han-Seong Kim, Dong-eun Lee, Seog-yun Park, Chi Chiu Wang, Young-Ho Kim, Tae-Min Kim, Seong Gu Heo, Namshik Han, Kyeong-Man Hong

**Author notes:** These authors contributed equally to this work (Youngbeen Moon, Chung Hwan Hong). (Y.M.); (J-K.K.), (C.H.H.); (E-K.K.); (H.W.C.), (D-W.H.) (J-H.G.), (H-S.K.), (S-y.P.) (C.C.W.), (Y.-H.K.), (N.H.).

## Abstract

**Background:** We previously developed a benchmarking strategy using mixtures of homozygote and heterozygote DNAs as reference standards to simultaneously assess sensitivity and false positive (FP) error rates in targeted next-generation sequencing (T-NGS) and whole-exome sequencing (WES), revealing substantial variability across commercial platforms. However, optimal analytic conditions for clinical application remain undefined.

**Methods:** We systematically evaluated multiple sequencing kits and bioinformatics pipelines across various variant allele fraction (VAF) thresholds to identify conditions that maximize both sensitivity and specificity. Recurrent error-prone alleles were defined and filtered to enhance specificity.

**Results:** Optimal performance was achieved using the DRAGEN pipeline with recurrent FP allele filtering. For T-NGS, a 1% VAF cutoff yielded a 95% detection threshold of 2.99% and 1.21 FPs per megabase (FP/Mb); for WES, a 2% cutoff yielded a 95% threshold of 5.02% and 1.15 FP/Mb. These settings improved sensitivity >3-fold and reduced FP rates >96% versus suboptimal pipelines. Notably, VAF thresholds flattened sensitivity differences across platforms, obscuring key performance disparities—challenging assumptions that T-NGS is inherently more sensitive than WES. In-house and conventional pipelines undercalled up to 10% of true variants. Restricting reporting of 1–4% VAF variants to ∼1,000 predefined actionable sites enabled recovery of clinically relevant mutations while reducing FP risk >99%.

**Conclusions:** This study provides a quantitative framework for optimizing NGS performance. Our findings support actionable strategies to improve diagnostic accuracy in clinical genomics through tailored pipeline selection, VAF thresholding, and artifact filtering.

## Background

Next-generation sequencing (NGS) enables high-throughput detection of DNA variants and is widely adopted in both clinical and research contexts [1–3]. Despite its transformative impact, challenges in analytical sensitivity and specificity persist. Large-scale initiatives such as SEQC2 have promoted quality control by developing reference standards for whole-genome, whole-exome, and single-cell sequencing [4–12], influencing regulatory frameworks from the FDA and CLIA [13–15]. However, clinical laboratories are not required to assess test sensitivity or specificity, in part due to the absence of practical evaluation methods.

NGS error rates are often underestimated, with reliance on bioinformatics pipelines to correct sequencing artifacts [16–18]. Yet, discrepancies in whole-exome sequencing (WES) results—up to 43% between major cancer databases—highlight the prevalence of false negatives, particularly when compared to targeted NGS (T-NGS) [19,20]. Sensitivity assessments using pooled HapMap DNA are limited to predefined variants and narrow VAF ranges (<2.5%), restricting their generalizability [21].

To reduce false positives (FPs), clinical laboratories often apply stringent variant allele fraction (VAF) thresholds, such as a 5% cutoff, potentially excluding low-VAF mutations of clinical relevance [22,23]. Conversely, more permissive thresholds may increase FP rates. Existing specificity assessments typically rely on HapMap samples or formalin-fixed paraffin-embedded (FFPE) tissues with known mutations [24–28], neither of which provide comprehensive error profiles.

To address these limitations, we recently developed a flexible benchmarking strategy using reference-standard mixtures (RSMs) composed of homozygote (hydatidiform mole) and heterozygote (blood) DNA to concurrently evaluate sensitivity and FP error rates in both T-NGS and WES [29]. This approach revealed marked variation in detection thresholds (1.5–24.7% for T-NGS; 5–20% for WES) and FP rates (5.8 × 10^−4^ to 1.4 × 10^−7^ for T-NGS; 130 to >30,000 FPs/sample for WES), emphasizing the need for standardized, quantitative metrics.

Although sensitivity and specificity are typically assessed separately, they are inversely related: higher VAF cutoffs reduce FPs at the expense of sensitivity [29–31]. In this study, we systematically evaluate how VAF thresholds influence this trade-off across platforms and pipelines, and we propose strategies to improve the detection of low-VAF mutations while minimizing FP errors—offering practical solutions for improving the reliability of NGS in clinical settings

## Methods

### NGS data generation using reference-standard DNA mixtures

We utilized previously published datasets [29,31] generated using mixtures of homozygote (HoZ), heterozygote (HetZ), and mixture DNAs, with IRB approval from the National Cancer Center, Korea. Homozygous DNA was derived from a hydatidiform mole (NA07489, Coriell), and heterozygote DNA from human blood.

Targeted NGS (T-NGS) data were generated by four commercial providers using the following assays: Company AA (Macrogen, Seoul) with Axen™ Cancer Panel, Company BB (Theragen Bio, Seoul) with SureSelect™ Cancer CGP Assay, Company CC (Geninus, Seoul) with CancerSCAN™, and Company DD with TruSight™ Oncology 500 (TSO500, Illumina) which incorporated unique molecular identifiers (UMIs).

All T-NGS assays used 200 ng input DNA, except for TSO500 (30 ng). Whole-exome sequencing (WES) was performed by: Company AA with SureSelect™ Human All Exon v8 (Agilent), Company BB with SureSelect™ XT Human All Exon v5 (Agilent), and Company CC with Human Exome v2.0 (Twist Biosciences).

Each WES assay used 500 ng input DNA. DNA quality and quantity were assessed per provider-specific SOPs. Library preparation, target capture, and paired-end sequencing were conducted using Illumina platforms. Companies AA and CC are CAP-accredited; Company BB holds Korean FDA accreditation for T-NGS.

### Variant calling and alignment

All reads were aligned to the GRCh37 (hg19) reference genome. For benchmarking, DRAGEN (v4.2, Illumina) was used to re-analyze raw T-NGS and WES data. Company DD data were processed with Illumina DRAGEN TSO500 Analysis Software (v2.1.1). Companies AA, BB, and CC also provided in-house variant calls processed through their proprietary pipelines.

For T-NGS DRAGEN analysis, Company BB provided a BED file, while Companies AA, CC, and DD limited variant calls to exonic regions of target genes.

For conventional analysis, raw T-NGS data from Companies BB and CC were aligned using BWA-MEM (v0.7.19) with default parameters, and variants were called using GATK Mutect2 (default settings) as described previously [31].

### Sensitivity benchmarking and detection thresholds

Alleles were classified by VAF as follows: null (N, VAF = 0), He-ll (0 < VAF < 0.01), He-low (0.01 ≤ VAF < 0.1), heterozygous (He, 0.10–0.75), and homozygous (Ho, ≥0.90). Sensitivity-informative (or informative) allele pairs (N–Ho, Ho–N, N–He) were defined by expected presence/absence patterns between HetZ and HoZ. A site was considered a false negative (FN) if the expected variant was absent in the mixture sample. Variants with <10 supporting reads in both T-NGS and WES were excluded from analysis.

Detection sensitivity was quantified by Probit regression using a cumulative normal distribution to estimate the 95% Limit of Detection (LOD)—the VAF at which 95% of expected variants were detected.

For platform comparisons, WES variants were restricted to the T-NGS target gene regions of each company, generating AA-, BB-, and CC-limited variant sets. Sensitivity was then compared between each T-NGS and its corresponding WES-limited dataset (e.g., BB-TNGS vs. BB-limited WES).

### Allele pair enumeration across platforms

To assess variant counts across platforms, genotype-defining allele pairs (e.g., Ho–N, N–Ho, N–He, Ho–Ho, Ho–He) were enumerated in the AA-, BB-, and CC-limited sets. Detection rates were calculated relative to DRAGEN-analyzed T-NGS (dTNGS) or DRAGEN-analyzed CC-limited WES.

### False positive error classification and specificity assessment

False positive (FP) errors were defined as: (1) N–N variants in reference-standard mixture (RSM) samples; (2) Secondary variants at sites already harboring a primary mutation; (3) Variants unique to HoZ or HetZ; and (4) Low VAF non-linear (lVnL) variants lacking expected dilution response (<10% VAF).

This composite FP definition was adapted from [29,31]. Variants with read depth <100 (T-NGS) or <20 (WES) were excluded. FP rates were calculated as FP/Mb within AA-, BB-, and CC-limited target regions.

### Distribution of mutations by VAF in cancer cell lines

To assess VAF distributions in real-world samples, we analyzed T-NGS data from 35 solid tumor cell lines across 151 genes, based on two independent datasets [20]. Observed mutations were stratified by VAF: 40–80%, 20–40%, 10–20%, 5–10%, 2.5–5%, 1.25–2.5%, and <1.25%.

These data were used to evaluate the detectability and clinical relevance of low-VAF mutations and inform strategies for VAF thresholding.

## Results

### Evaluation of sensitivity and specificity of NGS using mixtures of homozygote and heterozygote DNA

As shown in Fig. 1A, T-NGS data from Companies AA–DD and WES data from Companies AA–CC were analyzed using mixtures of homozygote (HoZ) and heterozygote (HetZ) DNAs as reference standard materials (RSMs) [29,31]. Variant calling was performed using company owned in-house pipelines, the Dynamic Read Analysis for GENomics (DRAGEN) pipeline, and a conventional BWA + Mutect2 pipeline. For consistent comparisons, only SNVs within each company’s T-NGS panel were considered; HLA genes were excluded. Note that company identifiers in this study follow the naming convention used in our previous T-NGS report [31]; accordingly, Companies AA, BB, and CC in this study correspond to Companies BB, CC, and AA, respectively, in our earlier WES report [29]. This ensures consistency in platform-specific comparisons throughout the study.

**Figure 1.**
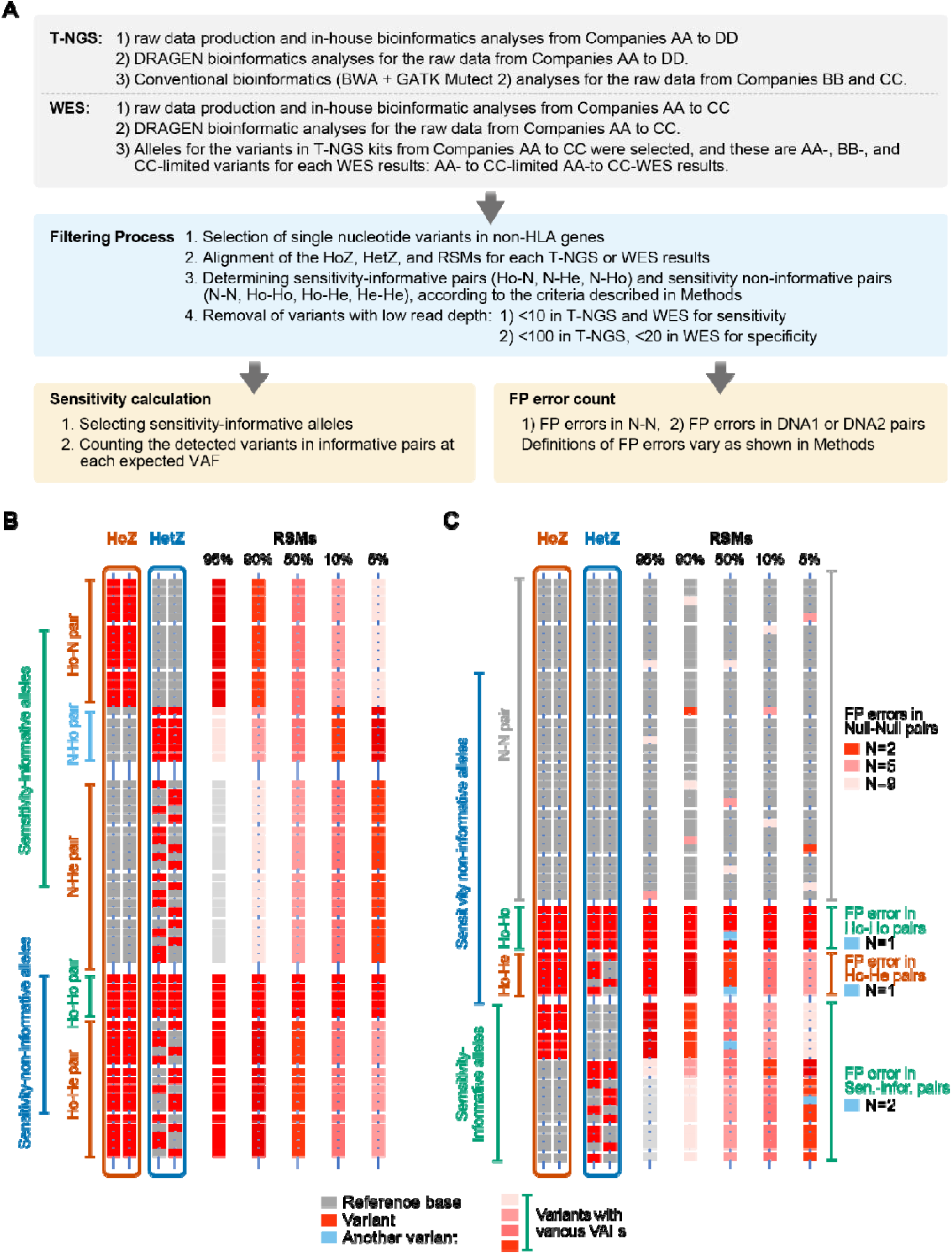
Study workflow and evaluation strategy for NGS sensitivity and specificity using DNA mixture standards. (A) Overall study design. Single-nucleotide variants (SNVs) in non-HLA genes were analyzed using four targeted NGS (T-NGS) kits (AA to DD) and three whole-exome sequencing (WES) kits (AA to CC) from four commercial providers. Sensitivity-informative (or informative) and non-informative alleles were classified to assess analytical sensitivity and specificity. **(B)** Strategy for sensitivity assessment using informative allele pairs from the allelic status of HoZ (H. mole) and HetZ (blood) DNAs. Reference bases and variants are color-coded, and the detection of informative alleles (e.g., N-Ho, Ho-N, N-He) across mixtures allows estimation of variant detection sensitivity. **(C)** Strategy for false positive (FP) error evaluation. FP calls were quantified in target sequences including informative (N-Ho, Ho-N, N-He) and non-informative (Ho-Ho, Ho-He, He-Ho, N-N, etc.) pair alleles. For B and C, reference standard mixtures (RSMs) are various mixtures of HoZ and HetZ (e.g., 95:5 to 5:95).

Alleles were classified by VAF values into null (N, homozygous reference base), homozygous variant (Ho), and heterozygous variant (He). In addition, the allele definition extended to include He-low and He-ll alleles, both of which have VAF values less than 0.1. Allele pairs were categorized based on allele status in HoZ and HetZ DNAs; for example, Ho–N refers to an allele that is homozygous variant in the HoZ sample and null in the HetZ sample.

Sensitivity (Fig. 1B) was calculated from detection rates across expected variant allele fractions (eVAFs) using sensitivity-informative (or informative) allele pairs (Ho–N, N–He, N–Ho). Specificity (Fig. 1C) was assessed as false positive (FP) error rates per megabase, based on the presence of unexpected variants in target sequences including both non-informative (N–N, Ho–Ho, Ho–He) and informative (Ho–N, N–Ho, N–He) allele pairs, as well as allele pairs containing He-low and He-ll alleles.

### Effects of VAF cutoffs on sensitivity of T-NGS and WES

NGS optimization strategies often involve deeper sequencing to enhance sensitivity and the use of higher VAF cutoffs to reduce false positives (FPs) [32–34]. In clinical practice, variants below 5% VAF are frequently excluded. However, the systematic impact of such cutoffs on sensitivity has not been thoroughly evaluated.

As expected, applying a 1% VAF cutoff led to decreased sensitivity in most T-NGS datasets compared to the 0% cutoff (Figs. 2A, 2B). Interestingly, T-NGS data processed by in-house pipelines from Companies AA and BB showed no difference in sensitivity between the 0% and 1% cutoffs, likely due to stringent internal filtering criteria already embedded in their analysis pipelines.

**Figure 2.**
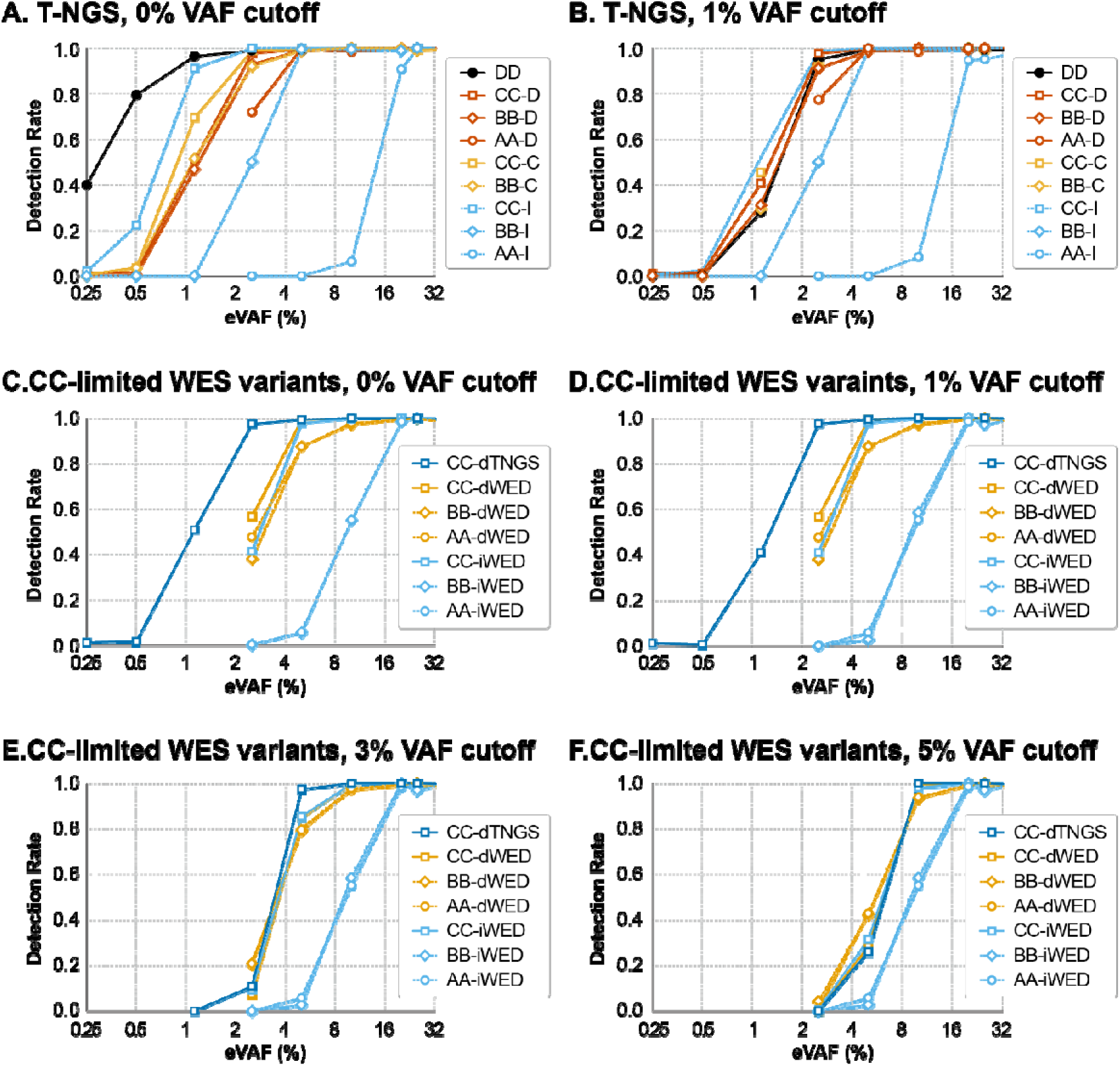
Relative sensitivity of T-NGS and WES results under varying VAF cutoffs. (A–B) Sensitivity of T-NGS data across all tested kits at variant allele frequency (VAF) cutoffs of 0% (A) and 1% (B). (C–F) Sensitivity of both T-NGS and WES for a subset of variants limited to the target regions of the T-NGS kit from Company CC (“CC-limited WES variants”) at VAF cutoffs of 0% (C), 1% (D), 3% (E), and 5% (F). In panels A and B, company identifiers (AA to DD) are followed by the analysis pipeline used: D for DRAGEN, I for in-house, and C for conventional analysis. In panels C to F, each label includes the company (AA to DD), analysis pipeline (d for DRAGEN, i for in-house), and sequencing method (T-NGS or WES). X-axis: Expected VAFs (%) calculated from dilution ratios of mixed reference-standard DNAs. Y-axis: Detection rate of variants (%), used to assess analytical sensitivity at each VAF level.

To compare T-NGS and WES performance, WES variants confined to the target region of the Company CC T-NGS panel (“CC-limited WES variants”) were analyzed. DRAGEN-based WES results from Company CC demonstrated higher sensitivity than those from Companies AA and BB at VAF cutoffs up to 1% (Figs. 2C, 2D), likely reflecting the superior capture efficiency of Company CC’s WES kit [29]. As expected, increasing the VAF cutoff from 1% to 5% led to reduced sensitivity in DRAGEN-based WES data (Figs. 2D–2F). Notably, in-house pipelines from Companies AA and BB consistently showed lower sensitivity (Fig. 2C), likely due to aggressive filtering strategies in their WES bioinformatics workflows [29].

### Sensitivity flattening across kits and platforms by VAF cutoffs

Surprisingly, applying a 1% VAF cutoff eliminated sensitivity differences among DRAGEN- and conventionally analyzed T-NGS results from Companies BB to DD (Fig. 2B), despite large variations in sequencing depth (∼1,000–4,000×). This finding challenges the common assumption that higher sequencing depth directly enhances sensitivity.

A similar flattening effect was observed in CC-limited WES data. While DRAGEN-analyzed WES from Company CC initially showed superior sensitivity over Companies AA and BB at 0% and 1% VAF cutoffs (Figs. 2C, 2D), this advantage disappeared at 3% and 5% cutoffs (Figs. 2E, 2F). Likewise, T-NGS from Company CC outperformed WES in sensitivity at low cutoffs (0–1%; Figs. 2C, 2D), but this benefit diminished at 3% and vanished entirely at 5% VAF (Fig. 2F), undermining the assumption that T-NGS is inherently more sensitive than WES.

### Additional observations on sensitivity flattening by VAF cutoffs or in-house filters

Sensitivity flattening by VAF cutoff was not limited to Company CC’s T-NGS targets. It was also observed in WES data restricted to T-NGS target sets from Companies BB and DD (BB- and DD-limited WES variants), with convergence occurring at the 3% VAF cutoff (Figs. S1A–S1D, S1G–S1I). This consistency across different target sets suggests that the flattening effect is not gene set–specific, but instead reflects internal consistency within T-NGS kits.

Further supporting this, DRAGEN-analyzed WES from Company CC showed similar sensitivity flattening by VAF cutoffs across all three target sets (BB, CC, and DD) (Fig. S2), indicating high internal consistency of the platform. Comparable patterns were observed for DRAGEN-analyzed WES from Companies AA and BB (data not shown).

In contrast, in-house WES pipelines from Companies AA and BB exhibited flat sensitivity across all VAF cutoffs up to 7.5% (Figs. S2D–S2F), suggesting that their filters preemptively excluded variants below this threshold. Consequently, sensitivity flattening in these pipelines likely reflects aggressive pre-filtering rather than a genuine effect of VAF cutoff.

### Optimizing sensitivity and specificity in NGS

Sensitivity and specificity varied substantially depending on bioinformatics pipelines, sequencing platforms (T-NGS vs. WES), and VAF cutoffs—consistent with our previous findings [29,31]. High sensitivity is essential for discovery applications (e.g., novel biomarker detection), whereas specificity is critical in clinical diagnostics, where confirmatory testing options are limited.

For Company CC’s T-NGS kit, the in-house pipeline showed higher sensitivity than DRAGEN at 0% VAF cutoff (Figs. 3A, 3B), but this advantage disappeared at 1% VAF (Fig. 2B). However, the in-house pipeline generated significantly higher false positive (FP) error rates (>100 FP/Mb at 5% VAF cutoff) compared to DRAGEN (<10 FP/Mb) (Figs. 3E, 3F). Since DRAGEN’s sensitivity remained stable between 0% and 1% (Fig. 3A), DRAGEN with a 1% VAF cutoff offers the optimal balance for T-NGS in Company CC.

**Figure 3.**
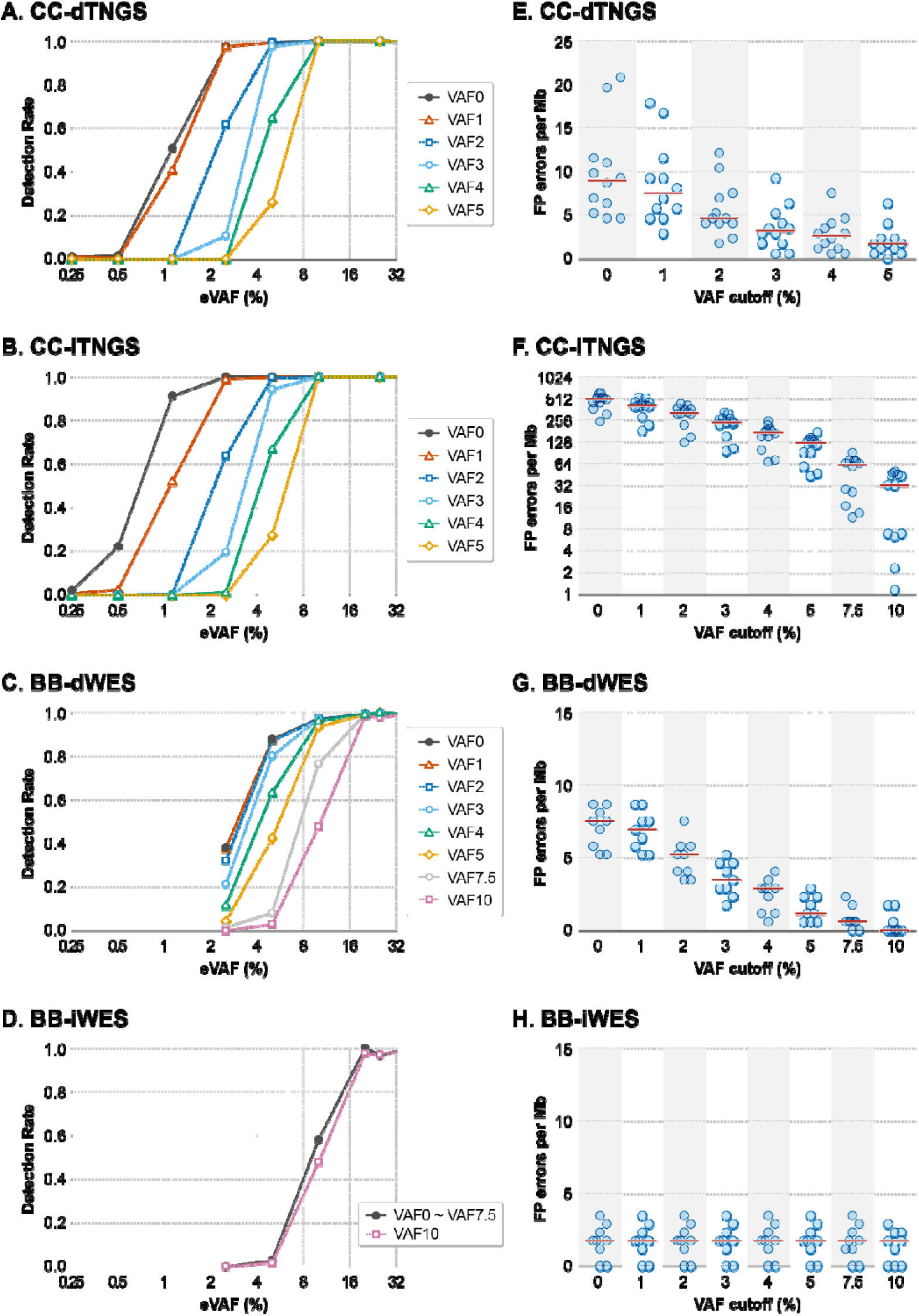
Impact of bioinformatics pipelines and VAF cutoffs on sensitivity and FP errors in T-NGS and WES. (A–B) Sensitivity of T-NGS results from Company CC analyzed using the DRAGEN pipeline (A) or in-house method (B). (C–D) Sensitivity of WES results from Company BB, limited to target variants from the T-NGS panel of Company CC, analyzed with DRAGEN (C) or in-house method (D). (E–F) FP error rates in T-NGS results from Company CC analyzed by DRAGEN (E) or in-house pipeline (F). (G–H) FP error rates in WES results from Company BB, using target variants restricted to the Company CC T-NGS panel, analyzed by DRAGEN (G) or in-house pipeline (H). For panels A–D, the X-axis represents expected variant allele frequency (VAF, %), and the Y-axis shows the detection rate (%) of sensitivity-informative variants. For panels E–H, the X-axis indicates the VAF cutoff applied (%), and the Y-axis represents the number of FP errors per megabase. In all panel labels (A–D), the suffix “VAF#” indicates the applied VAF cutoff (e.g., “VAF1” denotes a 1% cutoff).

For Company BB’s T-NGS kit, both DRAGEN and in-house pipelines produced low FP rates (<10 FP/Mb), but DRAGEN achieved higher sensitivity at VAF cutoffs up to 2% (Fig. S3). While the conventional pipeline matched DRAGEN in sensitivity, it showed more than threefold higher FP error rates (Figs. S3C, S3D, S3H, S3I), reinforcing the recommendation of DRAGEN with a 1% cutoff.

For Company DD’s T-NGS, the highest sensitivity was observed at 0% VAF, but this came with extremely high FP errors (2,776.8 FP/Mb; Figs. S3E, S3J). A 1–2% VAF cutoff under DRAGEN analysis significantly reduced FP errors while preserving sensitivity, indicating this as the optimal setting.

Taken together, across T-NGS kits from Companies BB to DD, the optimal analytic condition is DRAGEN analysis with a 1% VAF cutoff, offering consistently high sensitivity and minimal FP burden.

In WES, DRAGEN-analyzed results from all companies showed similar sensitivity at cutoffs up to 2% (Figs. 2C, 2D, 3C, S1), although FP error rates varied with VAF thresholds (Fig. 3G). In contrast, in-house WES pipelines from Companies AA and BB were largely insensitive to VAF cutoffs up to 7.5%, with low FP rates (<2 FP/Mb; Fig. 3H), but likely undercalled true low-VAF variants due to overly aggressive filtering (Fig. 3D). Therefore, DRAGEN analysis with a 2% VAF cutoff provides the best trade-off between sensitivity and specificity for WES.

### Risk of undercalling by in-house or conventional bioinformatics pipelines

WES variants restricted to T-NGS target regions (i.e., BB-, CC-, and DD-limited WES sets) showed consistent counts of genotype-defining allele pairs—Ho-N, N-He, N-Ho, Ho-He, and Ho-Ho—across DRAGEN-analyzed datasets (Fig. S4), despite the absence of BED files for CC and DD T-NGS panels. This consistency supports the reliability of cross-platform comparisons in sensitivity and specificity.

In contrast, in-house and conventional pipelines exhibited substantial undercalling. For example, Company BB’s in-house WES missed 7.21–8.01% of genotype-defining variants detected by DRAGEN across all three target sets (Fig. 4). Similarly, Company CC’s conventional pipeline missed approximately 10% of genotype-defining variants, recovering only 88.73% (T-NGS) and 89.47% (WES) of variants detected by DRAGEN. These findings highlight the risk of undercalling with in-house or conventional pipelines.

**Figure 4.**
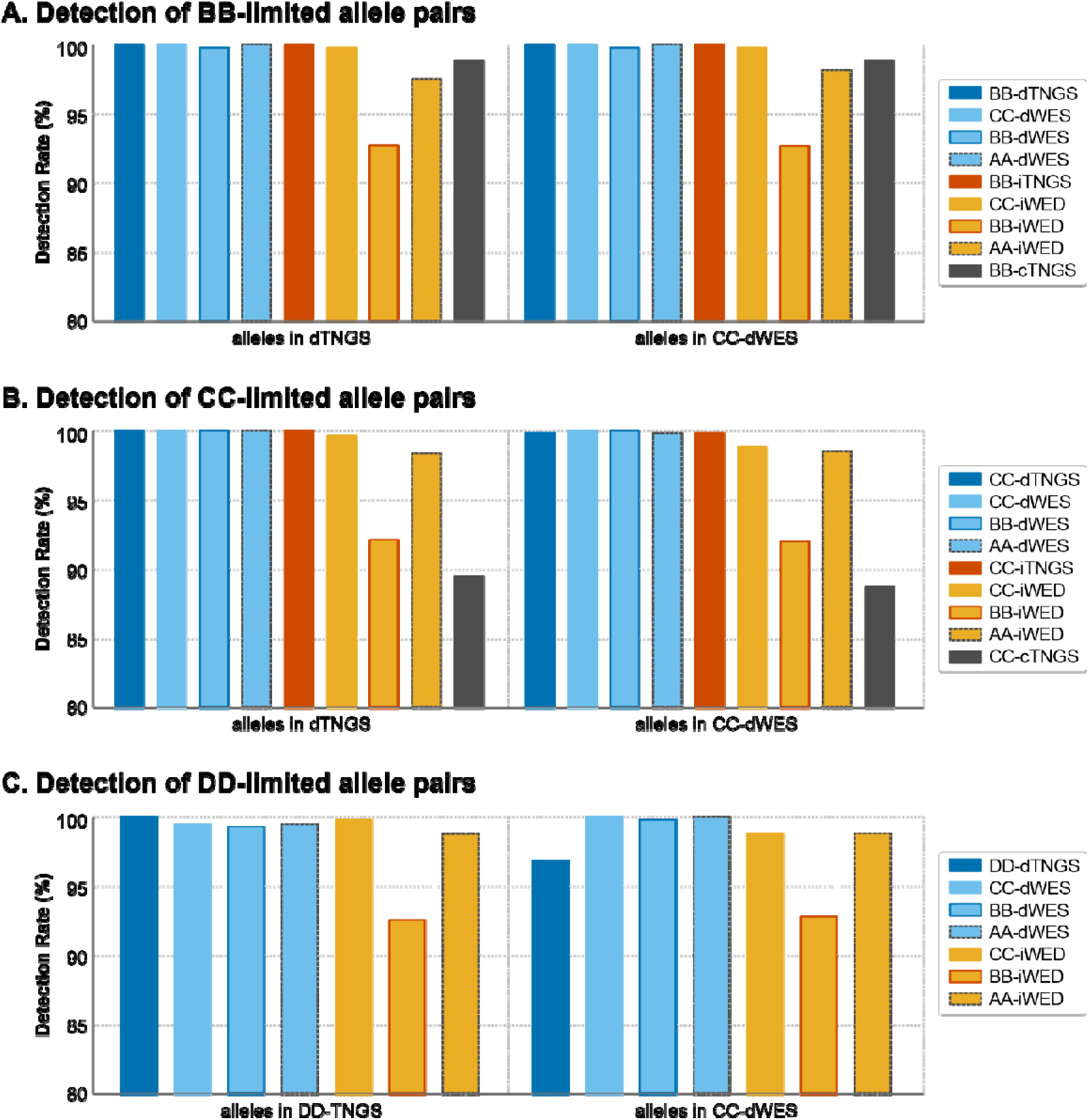
Difference in detection rate of allele pairs by various platforms, kits, and pipelines. The number of five allele pairs of Ho-N, N-He, N-Ho, Ho-He, and Ho-Ho were compared for variants limited to the target regions of T-NGS kits from Company BB (A), Company CC (B), and Company DD (C). Y-axis: Percentage of identified allele pairs compared to either DRAGEN-analyzed T-NGS results (left side), or DRAGEN-analyzed WES results (right side). In the figure labels, AA-, BB-, CC-, and DD-indicate the company responsible for raw data generation or kits. The prefix “i” denotes in-house analysis, “d” denotes DRAGEN-based analysis, and “c” denotes conventional analysis. TNGS and WES specify the sequencing platform used.

### Cross-platform comparison of FP error rates

This study employed a modified definition of FP errors (see *Methods*), incorporating low VAF non-linear (lVnL) artifacts—variants with <10% VAF that lack linear VAF correlations relative to dilution ratios. These lVnL artifacts often recur across HoZ, HetZ, and RSMs, and while some appear in N–N allele pairs, they are also present in non-N–N combinations.

FP error rates in T-NGS varied considerably (Fig. S4A). Under DRAGEN analysis at a 1% VAF cutoff, FP rates converged to ∼10 FP/Mb across Companies BB, CC, and DD (Fig. S5A): Company BB, 7.50 FP/Mb, Company CC, 3.62 FP/Mb, and Company DD, 17.53 FP/Mb.

By contrast, in-house and conventional pipelines showed markedly higher FP rates: Company CC in-house, 497.69 FP/Mb at 0% VAF, 124.06 FP/Mb at 5%; BB/CC conventional pipelines, >30 FP/Mb at 0%, remaining >10 FP/Mb at 5%; and Company DD in-house, extremely high FP rate of 2,776.8 FP/Mb at 0% VAF.

Similar trends were observed in WES. DRAGEN analysis with a 2% VAF cutoff reduced FP rates to <10 FP/Mb across BB-, CC-, and DD-limited WES sets (Figs. S5B–D). In contrast, Company CC’s in-house WES consistently produced FP rates of 20–38 FP/Mb across all cutoffs and target sets, further emphasizing the variability introduced by platform and pipeline differences.

### Reducing FP error rates via error-prone variant removal

We previously identified a set of FP error–prone alleles, primarily composed of recurrent low-VAF artifacts—referred to as low VAF non-linear (lVnL) variants—consistently detected across HoZ, HetZ, and RSM samples [31]. These variants can be systematically filtered without compromising sensitivity.

When variants at these error-prone allelic sites were removed, adjusted FP error rates were significantly reduced in both DRAGEN and in-house pipelines (*P* < 0.05, Mann–Whitney U; Fig. 5), demonstrating the pipeline-independent benefit of this approach. Specifically, adjusted FP rates decreased by 75.0–81.8% in DRAGEN-analyzed datasets and by 90.3–99.7% in in-house analyses, underscoring the effectiveness of recurrent artifact filtering in improving specificity (Fig. 5).

**Figure 5.**
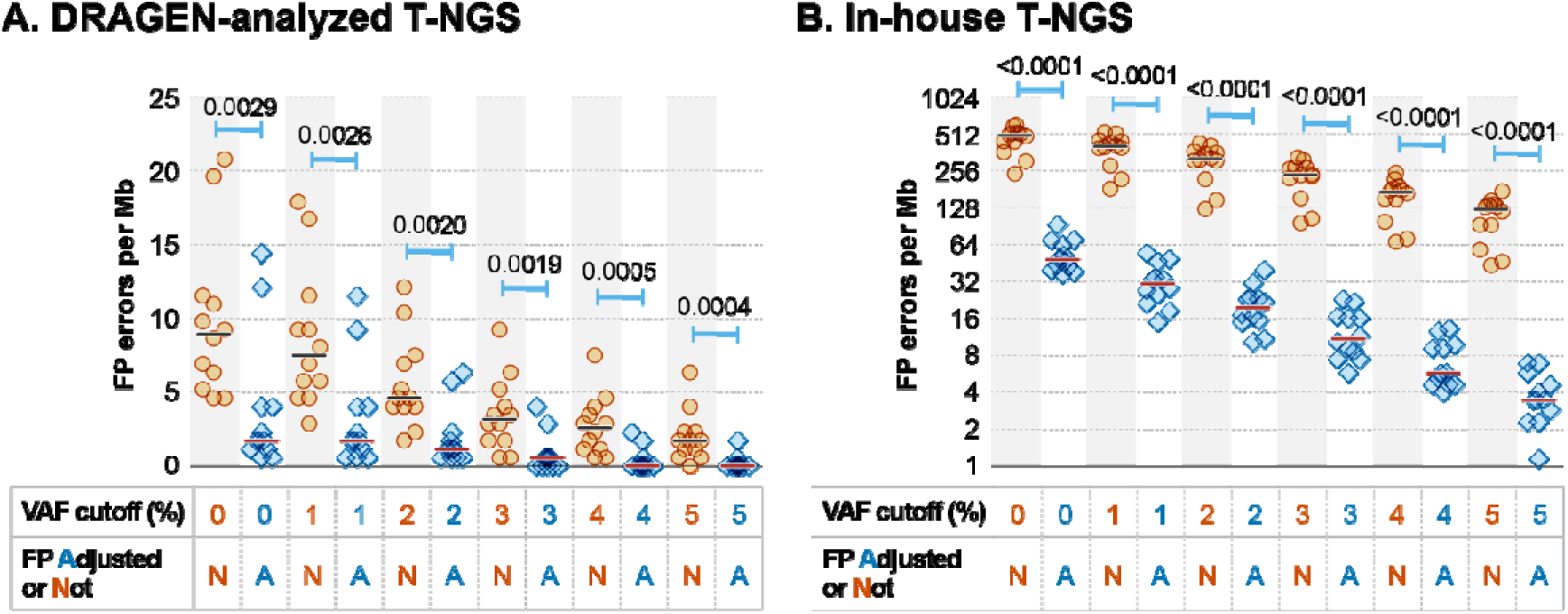
Effect of error-prone allele filtering on FP rates in Company CC T-NGS. (A) FP error rates in DRAGEN-analyzed T-NGS results from Company CC. (B) FP error rates in in-house T-NGS results from Company CC. P values for the difference in FP error rates between FP errors before (N) and after (Adjusted, or A) the removal of FP errors at error-prone allelic sites. X-axis, Grouped by VAF cutoff (%), and adjusted FP errors or not (Ad or not). Y-axis, Number of FP errors per megabase (Mb).

### Improvements in sensitivity and FP error rates under optimized conditions for T-NGS and WES

Under the optimal analytical configuration for T-NGS—DRAGEN analysis with a 1% VAF cutoff—Company CC (CC-dTNGS) achieved the highest sensitivity, with a 95% detection threshold of 2.99%, as estimated by Probit regression (Fig. 6A). In contrast, the poorest sensitivity was observed in Company BB’s in-house pipeline, with a 95% threshold of 9.68% at a 5% VAF cutoff, a value commonly used in clinical practice to minimize FP errors. This reflects a 3.24-fold difference in sensitivity between optimal and suboptimal configurations (9.68/2.99). Company AA’s T-NGS data were excluded from this comparison due to severely impaired sensitivity in its in-house pipeline.

**Figure 6.**
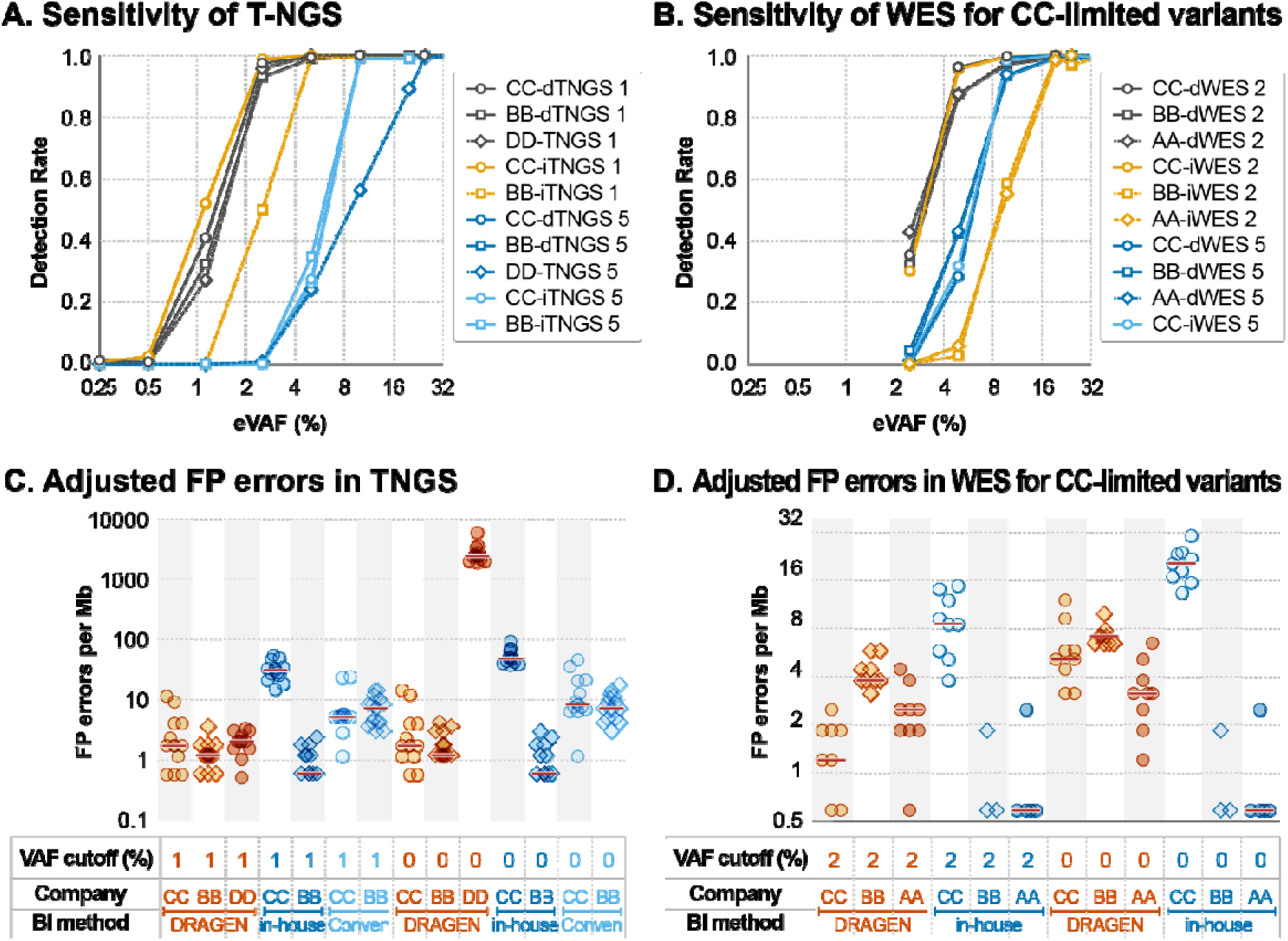
Improvements in sensitivity and FP rates under optimized conditions for T-NGS and WES. (A-B) Relative sensitivity improvements observed in DRAGEN-analyzed T-NGS (A; 1% VAF cutoff) and WES (B; 2% VAF cutoff) results, compared to those obtained under suboptimal conditions. Analyses were restricted to variants within the target regions of the T-NGS panel from Company CC. (C-D) Adjusted FP error rates for DRAGEN-analyzed T-NGS (C; 1% VAF cutoff) and WES (D; 2% VAF cutoff), shown alongside results from suboptimal pipelines. For both (B) and (D), analyses were limited to variants shared with the T-NGS panel from Company CC. X-axis (A–B): Expected variant allele frequency (VAF, %) based on predefined DNA mixture ratios. Y-axis (A–B): Detection rate (%) of sensitivity-informative alleles. X-axis (C–D): Grouped by VAF cutoff (%), data-generating company or kit (AA–DD), and bioinformatics pipeline (DRAGEN, in-house, or conventional). Y-axis (C–D): Number of FP errors per megabase (Mb). In panels for A–B, labels include the originating company (AA–DD), the bioinformatics method (d: DRAGEN; i: in-house), and the sequencing platform (T-NGS or WES).

For WES variants restricted to the T-NGS target region of Company CC (CC-limited WES variants), the optimal performance was achieved using DRAGEN with a 2% VAF cutoff, yielding the lowest 95% detection threshold of 5.02% (Fig. 6B). The poorest WES sensitivity was observed in Company BB’s in-house pipeline, with a 95% threshold of 19.51% across the 0–5% VAF range, representing a 3.89-fold difference (19.51/5.02). Similar differences between optimal and suboptimal WES conditions were also observed for BB- and DD-limited WES variants (Figs. S6A, S6B).

Following removal of recurrent error-prone variants (adjusted FP analysis), FP error rates in DRAGEN-analyzed CC-limited T-NGS at 1% VAF were substantially reduced to 1.21, 1.73, and 2.06 FP/Mb for Companies BB, CC, and DD, respectively (Fig. 6C). These adjusted rates were markedly lower than both the unadjusted FP rates (Fig. S5A) and the in-house CC T-NGS adjusted rates at 0–1% VAF (Fig. 6C). Although Company BB’s in-house T-NGS yielded the lowest adjusted FP rate (1.21 FP/Mb), its 95% detection threshold was 3.89-fold worse, indicating that the apparent specificity gain was driven by excessively stringent filtering. Thus, the best overall specificity for T-NGS was achieved by DRAGEN analysis of Company BB’s data (1.21 FP/Mb) without sacrificing sensitivity.

The worst specificity in T-NGS was observed in the unadjusted Company DD result at 0% VAF cutoff (2,776.80 FP/Mb; Fig. S5A), followed by the in-house CC pipeline (503.75 and 410.85 FP/Mb at 0% and 1%, respectively; Fig. S5A). Therefore, transitioning from these suboptimal conditions to the optimal configuration reduced FP rates by 99.96% (1.21 vs. 2,776.8) for DD and 99.86% (1.21 vs. 503.76) for in-house CC. Even compared to conventional CC analysis, the reduction was 96.57% (1.21 vs. 35.2).

For CC-limited WES, adjusted FP error rates under DRAGEN with a 2% VAF cutoff were 2.31, 3.46, and 1.15 FP/Mb for Companies AA, BB, and CC, respectively (Fig. 6D)—significantly lower than the unadjusted rates of 2.89, 5.19, and 2.88 FP/Mb (Fig. S5C). The worst WES specificity was observed in Company CC’s in-house WES, with 33.47 FP/Mb at 0% and 23.08 FP/Mb at 2% VAF. Thus, the optimal DRAGEN-based configuration reduced FP errors by 96.56% (1.15 vs. 33.47). Similar trends were seen for BB- and DD-limited WES (Figs. S5B, S5D, S6B, S6D), reinforcing the benefit of optimized pipelines and cutoff selection.

### Strategy to enhance NGS sensitivity

While stringent VAF cutoffs are effective at reducing false positive (FP) calls, they also eliminate many true low-VAF variants. Lowering these thresholds offers the potential to recover more clinically relevant mutations, with the magnitude of gain depending on the underlying VAF distribution—classified as decreasing, constant, or increasing models. Among these, the *increasing* model shows the greatest benefit from applying lower cutoffs (Figs. 7A, 7B).

**Figure 7.**
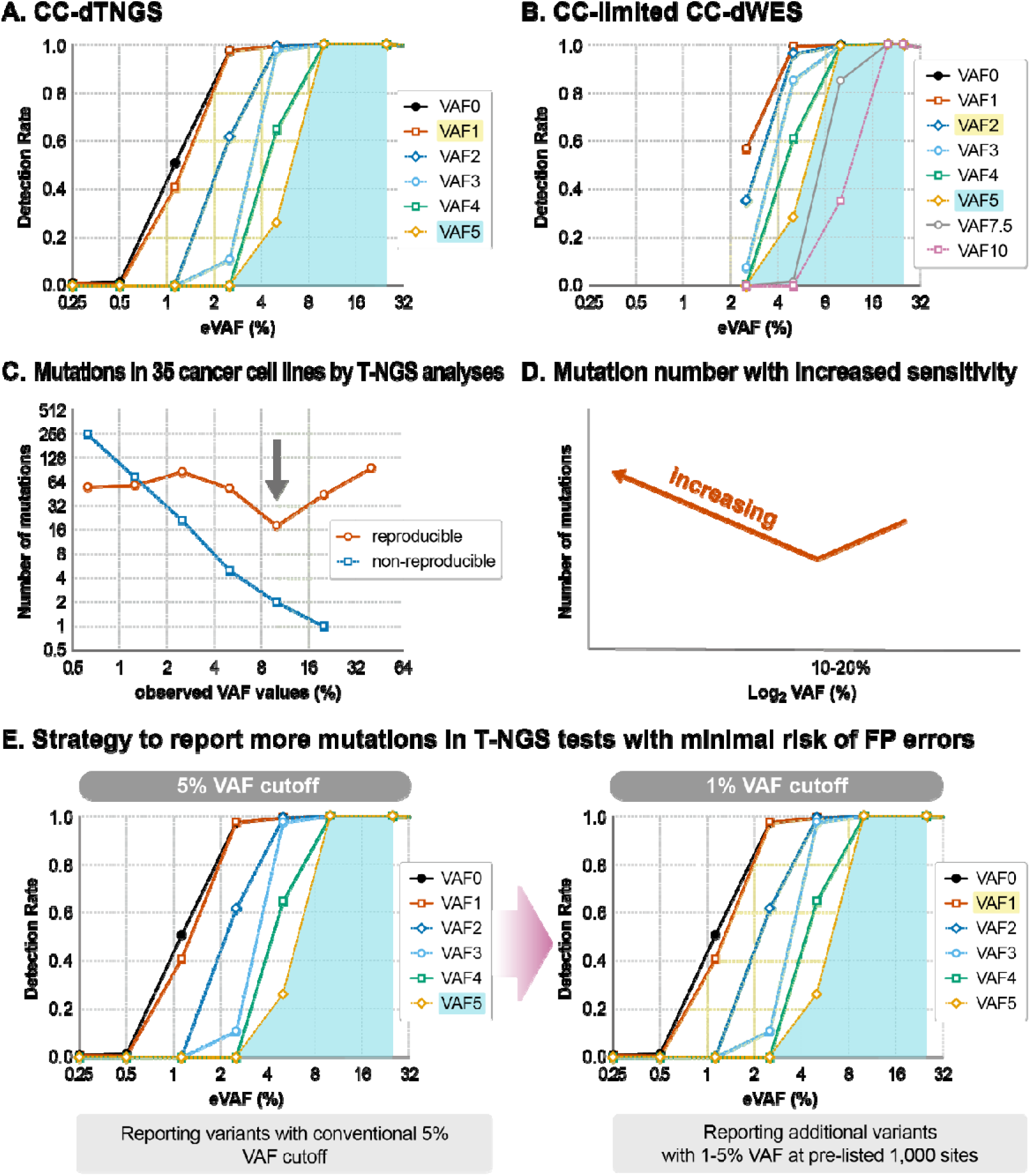
Sensitivity optimization via lower VAF cutoffs and mutation lists to reduce FP risk. (A–B) Enhanced variant detection with reduced VAF thresholds in DRAGEN-analyzed results from Company CC: T-NGS (A; 5% to 1% cutoff) and WES (B; 5% to 2% cutoff). Yellow shading indicates variants additionally detected at lower cutoffs. (C) Variant detection across varying VAF cutoffs in cancer cell lines using T-NGS. Orange indicates consistent variants detected in two independent raw datasets; blue indicates inconsistent calls. Green shading highlights a dip in mutation counts at 10–20% expected VAFs (gray arrow). (D) Revised model for interpreting mutation detection trends with decreasing VAF cutoffs, accounting for the result from cancer cell lines. (E) Schematic of a dual strategy to report more mutations while minimizing FP errors: Left: traditional 5% VAF cutoff reporting strategy. Right: enhanced reporting of lower-VAF variants (1–5%) restricted to a predefined list of clinically relevant mutations at 1,000 sites to avoid FP inflation. X-axes, VAF (%) or expected VAF values. Y-axes, Variant counts or detection rates (%).

To evaluate this effect in actual cancer samples, we reanalyzed targeted NGS (T-NGS) data from 35 solid tumor cell lines across 151 gene targets [20], using two independent datasets to assess mutation call consistency across VAF values. We observed a dip in consistent mutation calls at 10–20% VAF, followed by a rise as VAF decreased to ∼2.5%. Below ∼2.5%, consistent calls plateaued while inconsistent (noise) calls increased, indicating a limit of detection (Fig. 7C). Overall, the number of detectable mutations increased progressively as the VAF threshold was halved, highlighting that lenient VAF cutoffs (<5%) can substantially enhance mutation detection (Fig. 7D).

In clinical settings, a 5% VAF cutoff is frequently used to reduce FP errors, but this practice excludes a significant fraction of clinically actionable mutations. To balance sensitivity and specificity, we propose a hybrid approach: report all mutations >5% VAF, and for variants in the 1–5% range, report only those located within a predefined list of ∼1,000 actionable or high-value sites. Given the approximate 1.7 million base pair target space in Company CC’s T-NGS panel, this selective reporting strategy could reduce the probability of FP errors by ∼99.9% (1,000 / 1.7 × 10^6^), while preserving a large portion of clinically significant low-VAF mutations (Fig. 7E).

### Wobbling of observed VAF values

Observed VAFs showed considerable variability even at identical expected VAFs. For instance, variants expected to have a 10% VAF exhibited observed values ranging from approximately 5% to 15%—a two-fold variation. This degree of wobble was consistently observed across DRAGEN-analyzed T-NGS datasets from all three companies (Figs. S7A–S7C).

Such variability highlights the technical imprecision inherent in low-VAF measurements. Notably, an observed 5% VAF may in fact correspond to a true VAF of ∼10%. This is particularly relevant in clinical contexts: for example, a tumor with ∼30% mutated cells in a triploid (3n) genomic state is expected to produce a VAF of ∼10%, which may wobble down to 5% due to technical variability.

These findings underscore that low-VAF calls cannot be dismissed solely based on absolute thresholds, as they may represent biologically and clinically meaningful mutations affected by stochastic variation in measurement.

## Discussion

We applied a recently established benchmarking strategy using reference-standard DNA mixtures to systematically evaluate sensitivity and specificity in targeted next-generation sequencing (T-NGS) and whole-exome sequencing (WES) across multiple platforms, pipelines, and variant allele fraction (VAF) thresholds [29,31]. A key observation was that sensitivity differences across commercial kits and bioinformatics pipelines were substantially reduced—*flattened*—when standard VAF cutoffs were applied, including between T-NGS and WES platforms. These results challenge the common assumption that T-NGS is inherently more sensitive than WES [1,35], and suggest that fixed VAF thresholds may obscure underlying performance disparities.

By jointly analyzing sensitivity and false positive (FP) error rates under varying analytic conditions, we identified optimized settings for each platform: DRAGEN analysis with a 1% VAF cutoff for T-NGS and a 2% cutoff for WES. These configurations improved sensitivity by more than threefold compared to suboptimal in-house pipelines while maintaining low FP rates. Importantly, up to ∼10% of genotype-defining true variants were missed by in-house or conventional pipelines compared to DRAGEN-based analysis. This substantial undercalling reflects pipeline-related sensitivity loss and emphasizes the need for rigorous quantitative benchmarking when selecting or validating clinical NGS workflows.

FP error rates varied by pipeline and VAF threshold. Under permissive conditions, FP burden increased substantially. To address this, we systematically filtered error-prone variants—defined as recurrent, non-genotype-defining sites lacking VAF-dose correlation across reference mixtures. These likely represent systematic sequencing or alignment artifacts. Their removal led to dramatic FP reductions (75.0–99.7%) across pipelines, with >99% and >96% reduction in T-NGS and WES, respectively, under optimized conditions. These results underscore the critical role of artifact filtering in achieving high specificity without sacrificing sensitivity.

The clinical utility of low-VAF variants is well documented [20,22,23]. Due to VAF measurement variability (“wobbling”), an observed 5% VAF may reflect a true VAF closer to 10%—potentially indicating mutations present in ∼30% of tumor cells, especially in triploid genomes. Our reanalysis of real tumor samples revealed that actionable mutations within the 1–5% VAF range are frequent and increasingly detected with more permissive thresholds. However, standard clinical pipelines often exclude these variants due to rigid cutoffs.

To address this tradeoff, we propose a clinically actionable reporting strategy: report all variants above 5% VAF, while limiting reporting of 1–5% VAF variants to a pre-specified list of ∼1,000 known or potentially actionable sites. Given the ∼1.7 Mb target space of a typical T-NGS panel, this approach reduces the theoretical FP probability for low-VAF sites by ∼99.9% while recovering many clinically relevant variants otherwise missed. While similar “hotspot-only” recommendations exist in guidelines such as ESMO [36], implementation remains heterogeneous. Our results provide a reproducible framework to guide this practice with empirical sensitivity/specificity benchmarks.

While methods such as unique molecular identifiers (UMIs) [37,38] and ultra-deep sequencing [39] are often used to enhance low-VAF detection, our findings suggest that target recovery efficiency may be a more significant limiting factor. The TSO500 kit, which uses liquid-phase capture, achieved the best 95% detection threshold (∼1.6%) despite requiring 5–10× less input DNA than other platforms. These results highlight the value of optimizing capture chemistry in addition to sequencing depth or read de-duplication methods.

Beyond specific platforms, our benchmarking strategy offers a reproducible, platform-agnostic framework for evaluating pipeline performance. Open-source tools such as nf-core, Dockstore, and Galaxy [40,41] can incorporate these sensitivity and specificity benchmarks to improve transparency, reproducibility, and diagnostic validity. Moreover, these metrics may also serve as correction factors in AI-driven tools for variant interpretation [42,43], enabling more reliable inference in clinical genomics.

This study has limitations. All experiments were conducted using DNA from fresh blood or cultured cells, whereas most clinical samples are formalin-fixed, paraffin-embedded (FFPE) and subject to additional artifacts such as deamination-induced C>T changes [44,45]. We also excluded HLA genes due to alignment complexity [46], and did not evaluate small indels, which remain challenging due to low read support and alignment errors in cancer samples [47,48]. Extension of this framework to FFPE-derived DNA, HLA loci, and indels will be important for more comprehensive evaluation of clinical NGS performance.

## Conclusions

This study provides a quantitative framework for optimizing NGS-based clinical testing through the integration of variant allele fraction (VAF) thresholds, bioinformatics pipeline selection, and systematic filtering of recurrent error-prone variants. Applying these strategies enables clinical laboratories to simultaneously improve sensitivity and specificity, significantly reduce false positive burdens, and safely expand the reportable range to include clinically actionable low-VAF mutations. These refinements support more accurate and reliable genomic diagnostics, contributing to the advancement of precision medicine.

## Supporting information

supplementary informati

## Acknowledgements

The Bioinformatics Analysis Team assisted with DRAGEN-based raw data analysis; and the Biostatistics Collaboration Team supported the statistical analyses at the Research Core Center, National Cancer Center, Korea.

## Authors’ contributions

Conceptualization, K-M.H.; methodology, Y.M., W.C.H., J-K.K., E-K.K., H.W.C.; data acquisition and formal analysis, Y.M., W.C.H. J-K.K., D-W.H., J-H.K., H-S.K., S-y.P.; statistical analysis, D-e.L., Y.M., W.C.H.; data curation, Y.M., C.H.H.; writing-original draft preparation, Y.M., C.H.H., K-M.H.; writing-review and editing, C.C.W., Y-H.K., T-M.K., S.G.H., N.H.; supervision and funding acquisition, K-M.H.

## Availability of data and materials

All T-NGS fastq files used for this study are publicly available in the NCBI Sequence Read Archive (SRA) under the BioProject accession number PRJNA1134909. All WES fastq files used for this study are publicly available in the NCBI Sequence Read Archive (SRA) under the BioProject accession number PRJNA1178265.

## Ethics approval and consent to participate

The study received approval from the Institutional Review Board (IRB) of the National Cancer Center in Korea.

## Consent for publication

All authors have given their consent for publication.

