## supplementary informati for "Quantitative Optimization of Sensitivity and Specificity in Targeted and Whole-Exome Sequencing Using Reference-Standard DNA Mixtures"

**Table of contents**

**Supplementary figures S1–S7 and their legends**

Figure S1. Comparison of sensitivity between WES and T-NGS across varying VAF cutoffs.

Figure S2. Sensitivity of WES results at varying VAF cutoffs across different kits and pipelines.

Figure S3. Impact of VAF cutoffs on sensitivity and FP rates in BB and DD T-NGS results.

Figure S4. Consistency of sensitivity-informative and non-informative allele pair counts across T-NGS and WES platforms.

Figure S5. False positive (FP) error rates in T-NGS and WES results.

Figure S6. Optimized WES improves sensitivity and reduces FP errors for BB/DD T-NGS target regions.

Figure S7. Variability (“wobbling”) of observed VAF values in T-NGS results across expected VAF levels.

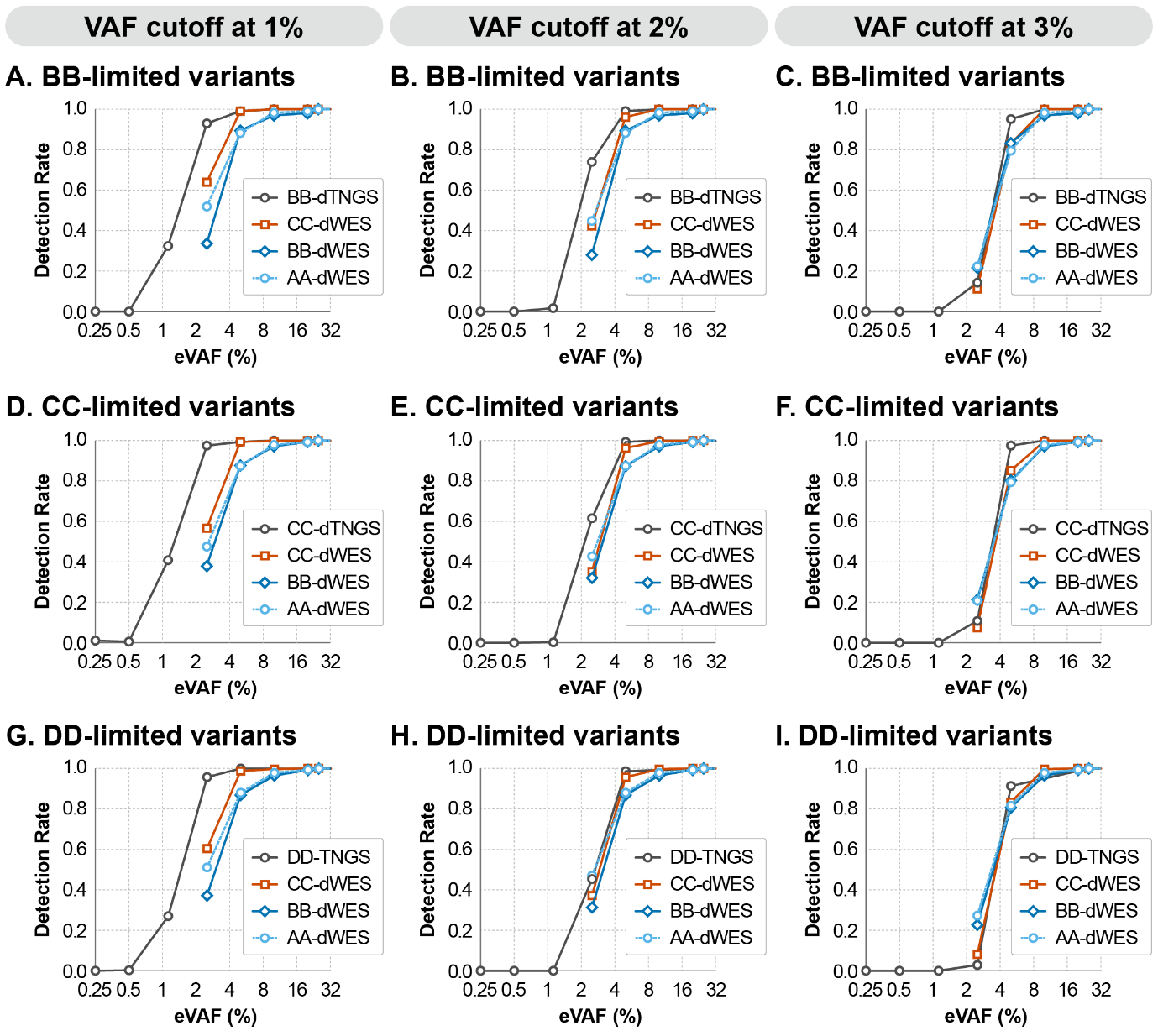

**Figure S1. Comparison of sensitivity between WES and T-NGS across varying VAF cutoffs.** (A–C) Sensitivity of DRAGEN-analyzed T-NGS data from Company BB (BB-dTNGS) compared to DRAGEN-analyzed WES data from Companies AA, BB, and CC (AA-dWES, BB-dWES, CC-dWES), using target variants restricted to the T-NGS panel of Company BB. VAF cutoffs: 1% (A), 2% (B), and 3% (C). (D–F) Sensitivity of DRAGEN-analyzed T-NGS data from Company CC (CC-dTNGS) and corresponding WES results from Companies AA, BB, and CC, using target variants restricted to the T-NGS panel of Company CC. VAF cutoffs: 1% (D), 2% (E), and 3% (F). (G–I) Sensitivity of DRAGEN-analyzed T-NGS data from Company DD (DD-dTNGS) and WES results from Companies AA, BB, and CC, using target variants restricted to the T-NGS panel of Company DD. VAF cutoffs: 1% (G), 2% (H), and 3% (I). Only data processed using the DRAGEN pipeline are shown.
X-axis: Expected VAF values (%) based on dilution ratios of reference-standard DNA mixtures. Y-axis: Detection rate (%) of sensitivity-informative variants.

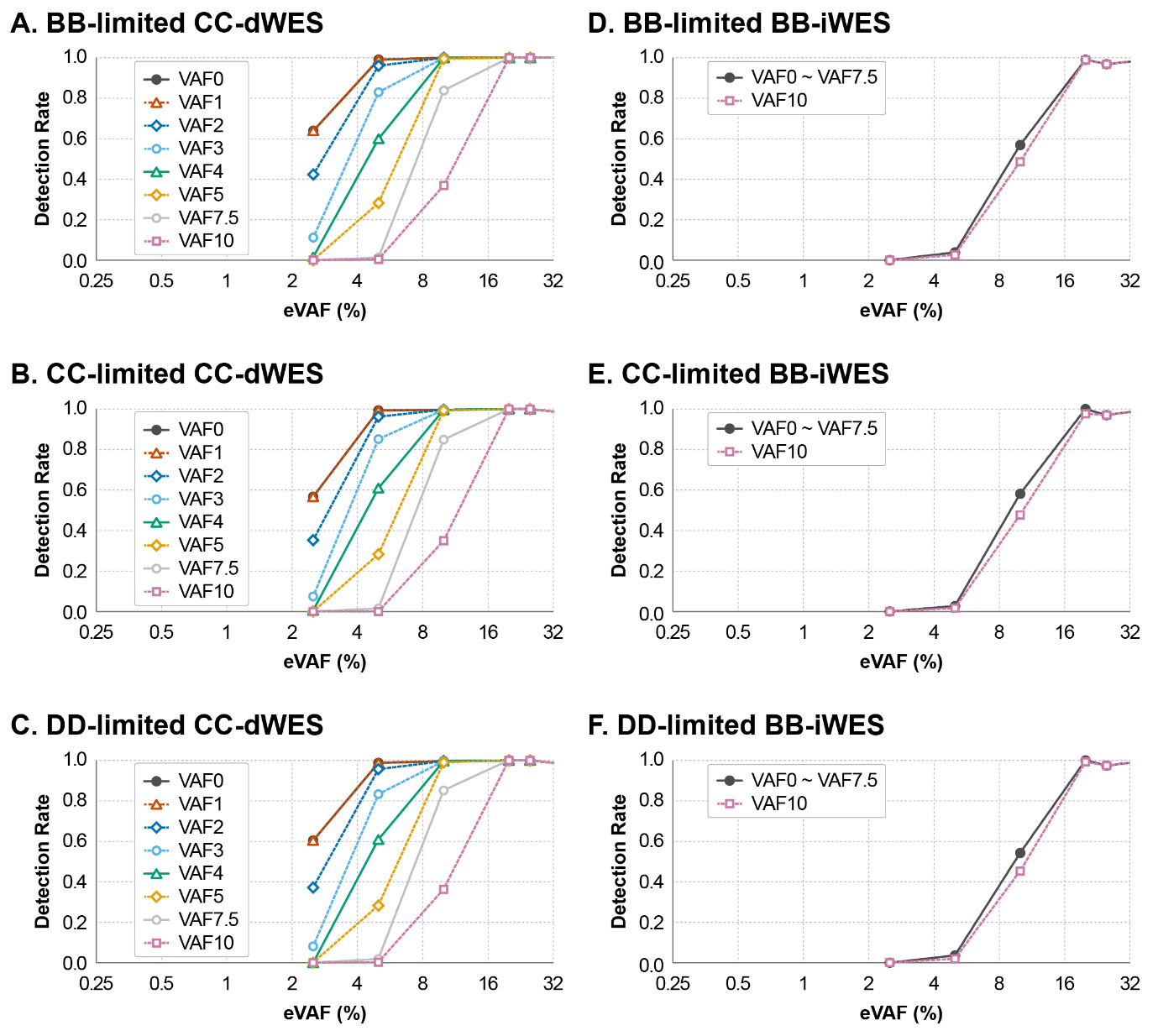

**Figure S2. Sensitivity of WES results at varying VAF cutoffs across different kits and pipelines.** (A–C) Sensitivity of WES results analyzed with the DRAGEN pipeline for variant sets limited to the target regions of the T-NGS kits from Company BB (A), CC (B), and DD (C). (D–F) Sensitivity of WES results analyzed with each provider’s in-house pipeline for the same variant sets from Company BB (D), CC (E), and DD (F). Each line represents a different VAF cutoff ranging from 0% to 10%, as indicated in the legend by the corresponding value followed by “VAF.” X-axis: Expected VAFs (%) derived from the known dilution ratios of the reference-standard DNA mixtures. Y-axis: Detection rate (% of expected variants detected), used to evaluate analytical sensitivity at each VAF level.

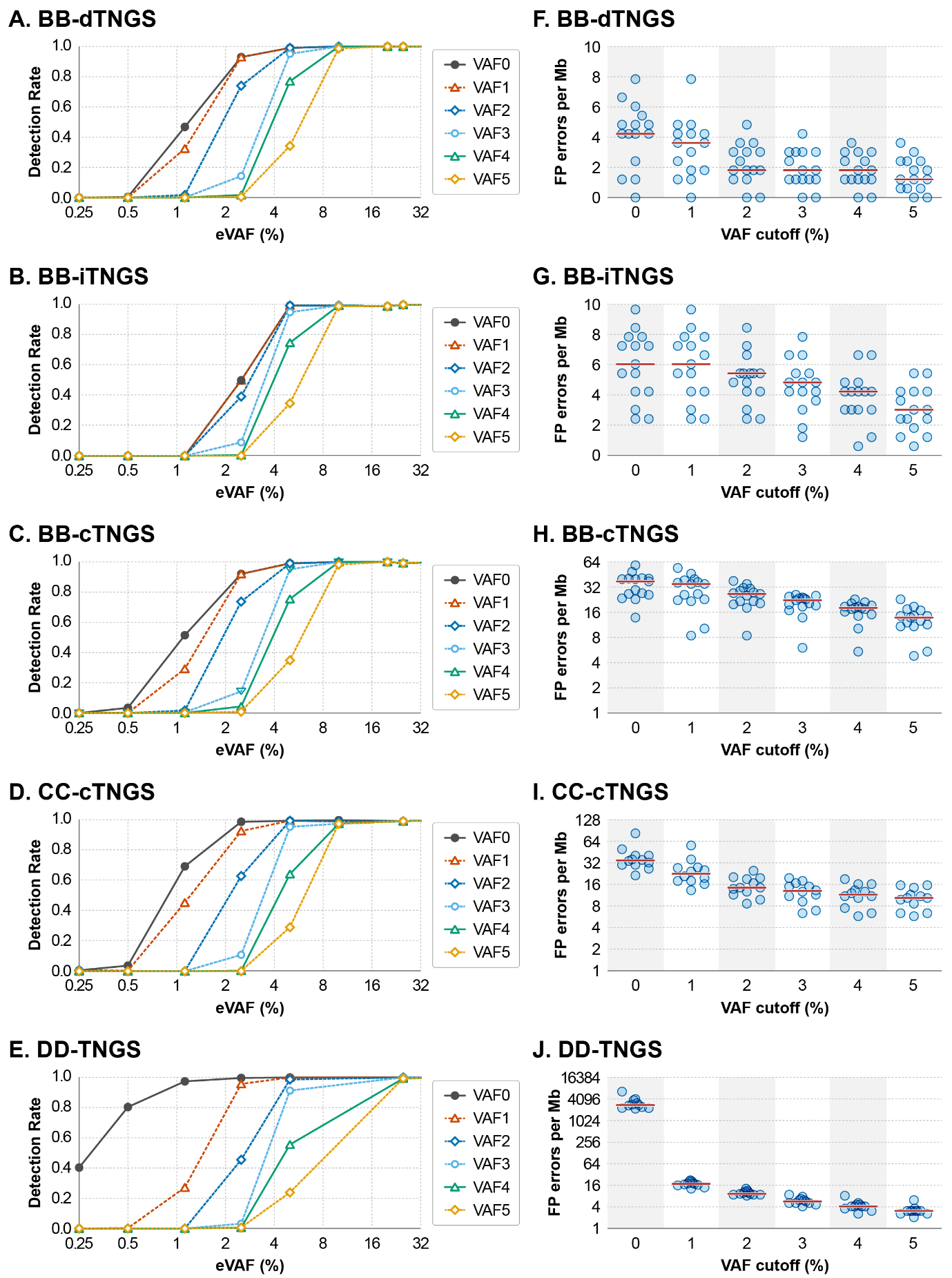

**Figure S3. Impact of VAF cutoffs on sensitivity and FP rates in BB and DD T-NGS results.** (A–C) Sensitivity of T-NGS results from Company BB analyzed using the DRAGEN (A), in-house (B), or conventional pipeline (C). (D-E) Sensitivity of T-NGS results from Company CC analyzed with conventional method (D), or Company DD analyzed with the DRAGEN pipeline (E). (F–H) FP error rates of T-NGS results from Company BB analyzed using DRAGEN (F), in-house (G), or conventional pipeline (H). (I-J) FP error rates of T-NGS results from Company CC analyzed with conventional method (I), or Company DD across varying VAF cutoffs (J). For panels A–E, the X-axis indicates expected variant allele frequency (VAF, %) derived from DNA mixture ratios, and the Y-axis represents the detection rate of sensitivity-informative variants. For panels F–J, the X-axis shows the applied VAF cutoff (%), and the Y-axis indicates the number of FP errors per megabase.

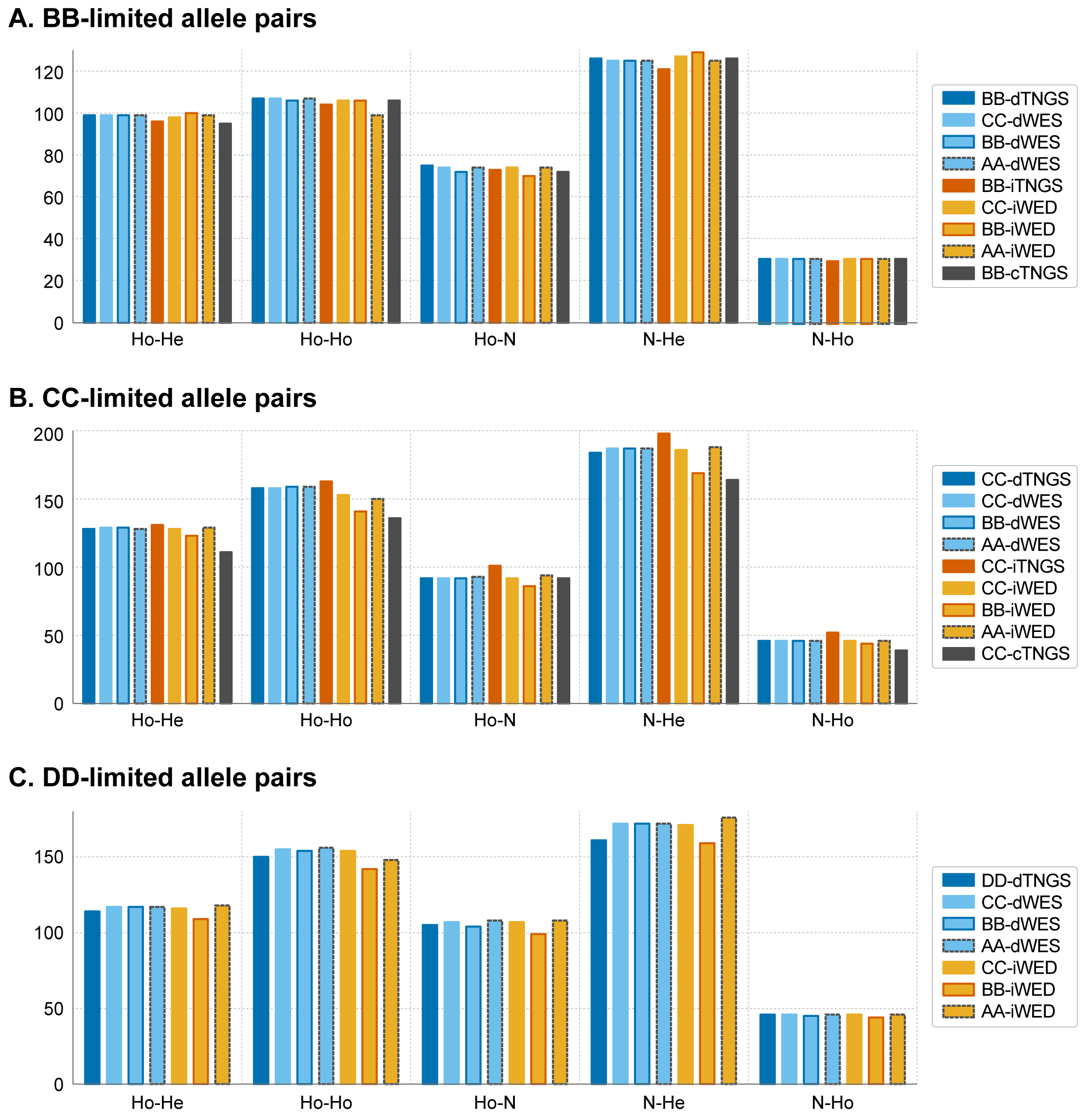

**Figure S4. Consistency of sensitivity-informative and non-informative allele pair counts across T-NGS and WES platforms.** Bar plots display the number of sensitivity-informative allele pairs (Ho-N, N-He, and N-Ho) and sensitivity-non-informative allele pairs (Ho-He and Ho-Ho) for variants limited to the target regions of T-NGS kits from Company BB (A), Company CC (B), and Company DD (C). X-axis: Type of allele pair. Y-axis: Number of allele pairs. In the figure labels, AA-, BB-, CC-, and DD- indicate the company responsible for raw data generation or kits. The prefix "i" denotes in-house analysis, "d" denotes DRAGEN-based analysis, and "c" denotes conventional analysis. TNGS and WES specify the sequencing platform used.

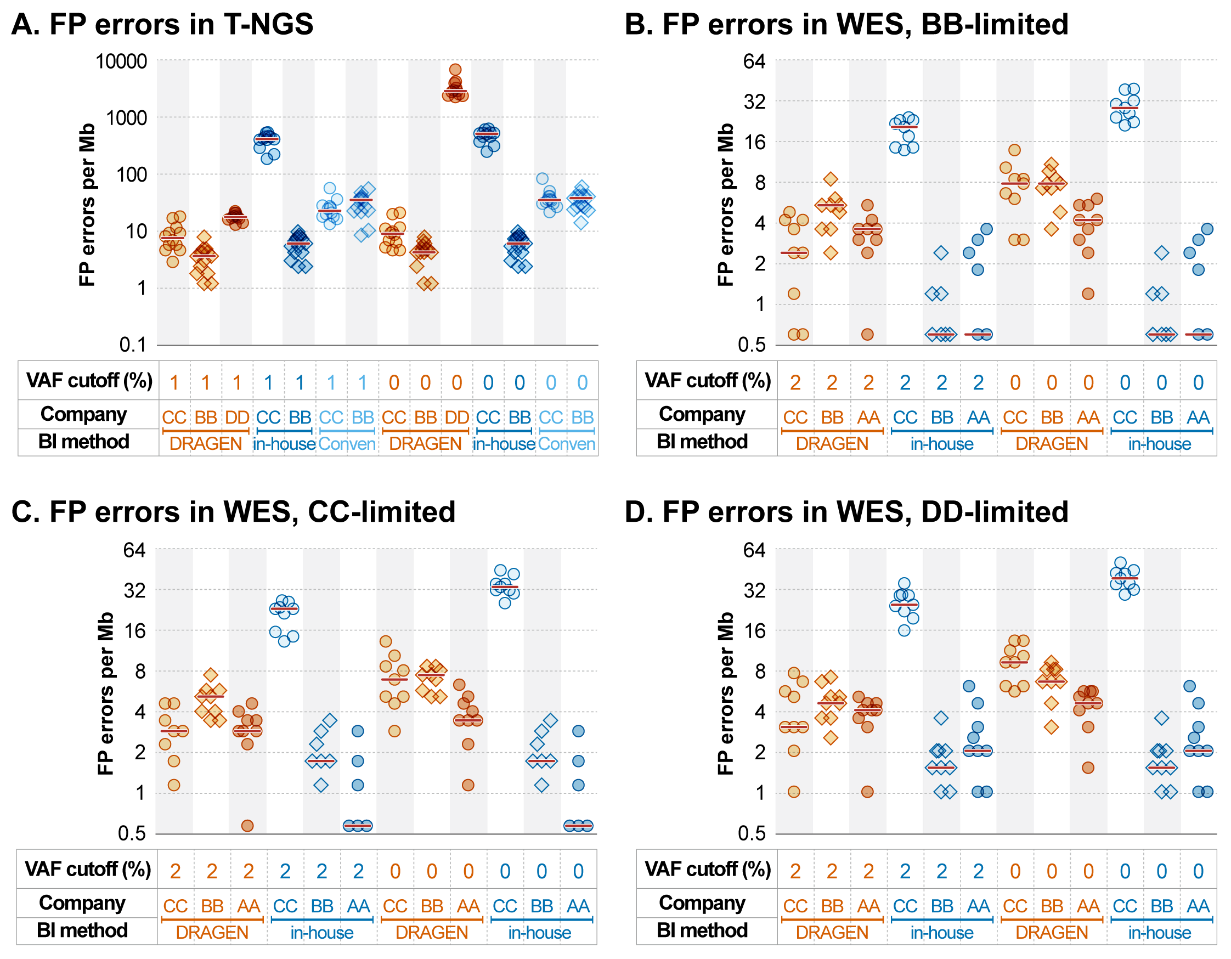

**Figure S5. False positive (FP) error rates in T-NGS and WES results.** (A) FP error rates in T-NGS results across different companies and analytical pipelines (DRAGEN or in-house) using VAF cutoffs of 0%, 1%, or 5%. (B–D) FP error rates in WES results for variants restricted to the target regions of T-NGS kits from Company BB (B), Company CC (C), and Company DD (D). X-axis: Results are grouped by VAF cutoff values (%), company identifiers (AA to DD), and bioinformatics methods (DRAGEN or in-house). Y-axis: Number of FP errors per megabase (Mb).

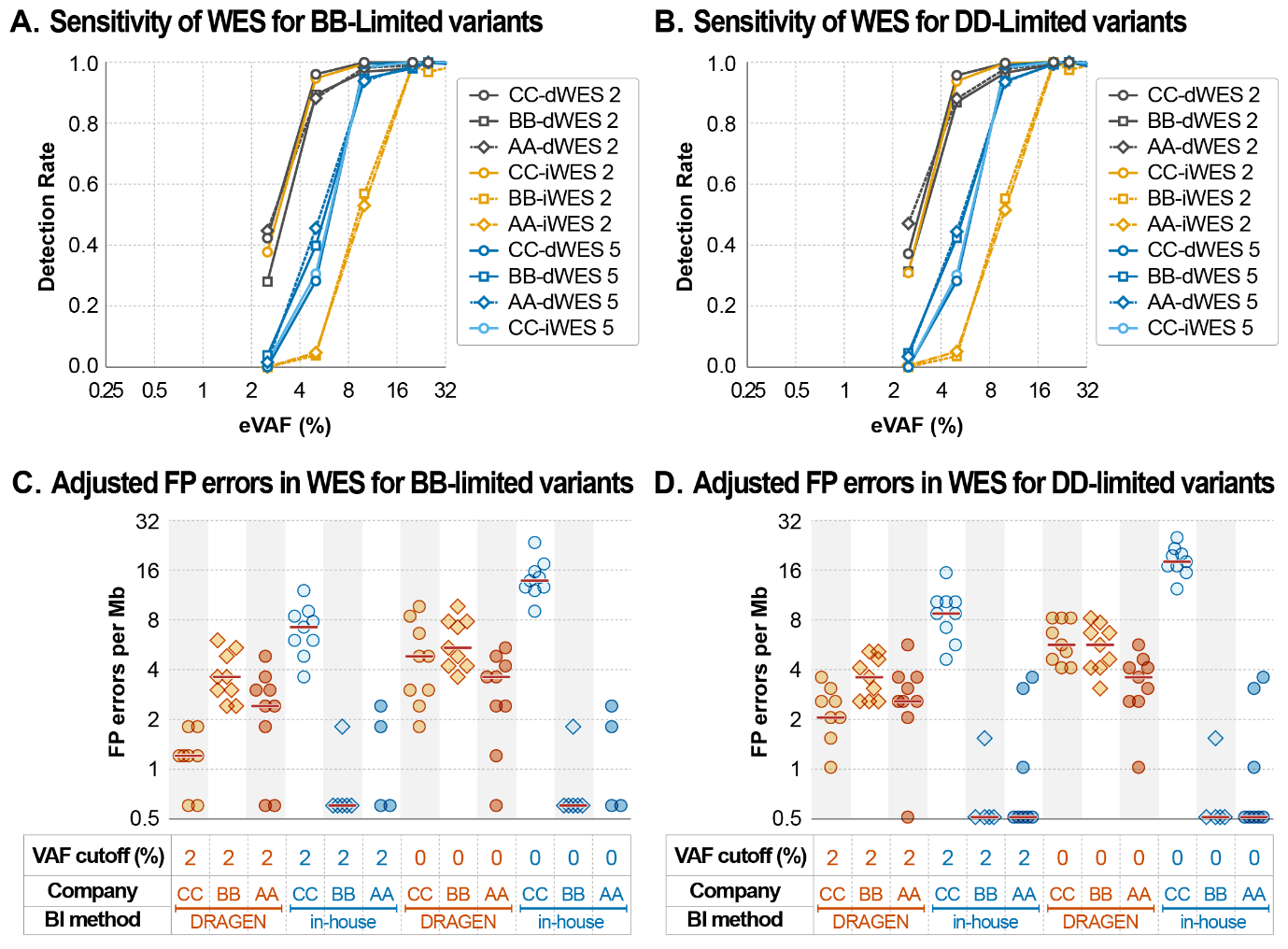

**Figure S6. Optimized WES improves sensitivity and reduces FP errors for BB/DD T-NGS target regions.** (A–B) Sensitivity of DRAGEN-analyzed WES results (2% VAF cutoff) compared to those obtained under suboptimal conditions for variants limited to the T-NGS target regions of Company BB (A) and Company DD (B). (C–D) Adjusted FP error rates in DRAGEN-analyzed WES results (2% VAF cutoff) compared to suboptimal conditions for the same target regions (C: BB-limited, D: DD-limited). Data points outside the axis limits (12 for C and 21 for D) are not shown. X-axis (A–B): Expected variant allele frequency (VAF, %) based on DNA mixture ratios. Y-axis (A–B): Detection rate (%) of sensitivity-informative variants. X-axis (C–D): Grouped by VAF cutoff (%), data-generating company (AA–DD), and bioinformatics pipeline (DRAGEN or in-house). Y-axis (C–D): Number of FP errors per megabase (Mb). In panel labels for A–B, AA-, BB-, and CC- represent the data-generating company, while prefixes "d" and "i" before TNGS or WES denote DRAGEN and in-house analysis, respectively.

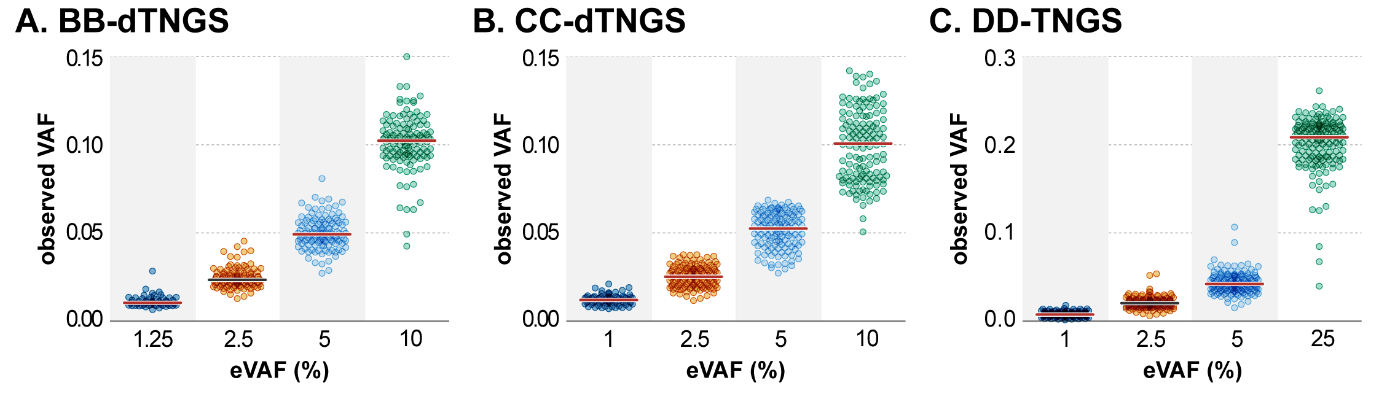

**Figure S7. Variability (“wobbling”) of observed VAF values in T-NGS results across expected VAF levels.** Scatter plots showing the fluctuation of observed variant allele fraction (VAF) values in DRAGEN-analyzed T-NGS results from Company BB (A), Company CC (B), and Company DD (C), plotted against the corresponding expected VAFs derived from reference-standard DNA mixtures. X-axis: Expected VAF values (%) based on known DNA mixing ratios. Y-axis: Observed VAF values (%) reported by the DRAGEN pipeline.
